# Central sensitization increases the pupil dilation elicited by mechanical pinprick stimulation

**DOI:** 10.1101/438788

**Authors:** E.N. van den Broeke, D.M. Hartgerink, J Butler, J Lambert, A Mouraux

## Abstract

High frequency electrical stimulation (HFS) of skin nociceptors triggers central sensitization, manifested as increased pinprick sensitivity of the skin surrounding the site at which HFS was applied. The aim of the present study was to compare the effects of HFS on pupil dilation and brain responses elicited by pinprick stimulation delivered in the area of increased pinprick sensitivity. In fourteen healthy volunteers HFS was applied to one of the two forearms. Before and twenty minutes after applying HFS, mechanical pinprick stimuli (64 mN and 96 mN) were delivered to the area surrounding the site at which HFS was applied as well as the contralateral control arm. During pinprick stimulation both the pupil size and electroencephalogram were recorded. HFS induced a clear and comparable increase in pinprick sensitivity for both the 64 and 96 mN stimulation intensity. Both pinprick stimulation intensities elicited a greater pupil dilation response when delivered to the area of increased pinprick sensitivity. However, this greater pupil dilation response was larger for the 64 mN compared to the 96 mN stimulation intensity. A similar pattern was observed for the negative wave of the pinprick-evoked brain potentials (PEPs), however, the increase was not significant for the 96 mN and showed only a trend towards significance for the 64 mN. These results show that there is a correspondence between the increase in pupil dilation and the increase in PEPs, but that pupil size is a more sensitive measure for detecting the effects of central sensitization than PEPs.

## INTRODUCTION

In humans, intense and/or sustained activation of skin nociceptors induces profound widespread increased mechanical pinprick sensitivity (a phenomenon typically referred to as “secondary hyperalgesia”, (Lewis, 1942; Hardy et al., 1945; LaMotte et al., 1991; Klein et al., 2004; van den Broeke et al., 2016). This widespread increased pinprick sensitivity is considered to be a manifestation of central sensitization (CS, Woolf, 2011; Baumann et al., 1991; LaMotte et al., 1991; Simone et al., 1991), which is defined by the International Association for the Study of Pain (IASP) as: *“increased responsiveness of nociceptive neurons in the central nervous system to their normal or subthreshold afferent input”* (Loeser and Treede, 2008).

To explore the changes in brain activity related to this increased pinprick sensitivity, we recently recorded pinprick-evoked brain potentials (PEPs), using two pinprick intensities (64 and 96 mN), before and after the induction of central sensitization using high-frequency electrical stimulation (HFS) of skin nociceptors (van den Broeke et al., 2017). HFS induced a significant increase in pinprick perception for both 64 mN and 96 mN pinprick stimuli delivered to the skin surrounding the area onto which HFS was applied. HFS also induced a significant increase in the magnitude of a late positive wave of PEPs (300-400 ms), maximal at the scalp vertex. However, contrary to the effect of HFS on pinprick perception, this increase was significant for the 64 mN pinprick stimulus but not for the 96 mN pinprick stimulus. A similar observation had been made in a previous study using capsaicin injection to induce increased pinprick sensitivity (van den Broeke et al., 2015). What could be the explanation for the fact that, unlike pinprick perception, the magnitude of this EEG response elicited by pinprick stimuli delivered to the area of secondary hyperalgesia is enhanced only for intermediate intensities of stimulation?

One possibility could be that the increase in magnitude of this positive EEG wave reflects an increase in cortical arousal mediated by the locus coeruleus (LC) - noradrenergic system (LC-NA). The LC is thought to play an important role in the regulation of arousal (Samuels and Szabadi, 2008; Sara and Bouret, 2012; Berridge, 2008) and its activity is increased following the presentation of a nociceptive stimulus (for review see Samuels and Szabadi, 2008).

LC neurons can exhibit two modes of activity; tonic and phasic. Tonic activity refers to the baseline rate of discharge of these neurons, whereas phasic activity refers to rapid and short-lasting increases in firing rate (Aston-Jones et al., 1999). Importantly, the occurrence of a phasic LC response appears to depend on the level of tonic/baseline activity. Indeed, Aston-Jones et al. (1999) showed in primates that periods of elevated tonic LC activity were consistently associated with decreased phasic responsiveness of LC neurons to target stimuli in a visual discrimination task. Furthermore, phasic LC responses to target stimuli were suppressed during drowsiness and very low LC tonic activity. Based on these observations it is hypothesized that when the level of arousal is either very high or very low, reflected in high or low levels of tonic activity, phasic LC responses are small or absent (Aston-Jones and Cohen, 2005).

If the processes underlying the increase in PEPs are at least partly mediated by phasic LC activity, one could hypothesize that the lack of increase of PEPs after HFS when participants are repeatedly exposed to high intensity pinprick stimuli (96 mN) as compared to when they are repeatedly exposed to stimuli of intermediate intensity (64 mN) is the result of a difference in baseline LC activity. The repeated application of high-intensity pinprick stimuli would increase baseline LC activity, and this would result in reduced pinprick-evoked phasic LC responses.

In primates, it has been shown that the activity of LC neurons is closely reflected in changes in pupil size, and that this is the case both for spontaneous and for stimulus-evoked activity (Joshi et al., 2006). Moreover, Reimer et al., (2016) showed in mice that rapid dilations of the pupil are tightly associated with phasic activity in noradrenergic axons in the cortex. Using pupil size as an index of LC activity, the aim of the present study was to test the hypothesis that the increase in PEPs following the experimental induction of CS is an electrophysiological correlate of LC-NA activity. More specifically, we studied the correspondence between the changes in PEPs and pupil size after HFS.

## METHODS AND MATERIALS

### Ethical approval

The experiment was conducted according to the declaration of Helsinki (except preregistration of the trial). Approval for the experiment was obtained from the local Ethical Committee (Commission d’Éthique Biomédicale Hospitalo-Facultaire) of the Université catholique de Louvain (UCL) (B403201316436). All participants signed an informed consent form and received financial compensation for their participation.

### Participants

Fourteen healthy volunteers took part in the experiment (7 men and 7 women; aged 19 – 28 years; 23.1 ± 2.9 years [mean ± sd]).

### Experimental design

The design of the experiment is summarized in Figure 1. Transcutaneous high frequency electrical stimulation (HFS) of the left or right volar forearm was used to induce increased pinprick sensitivity in the surrounding unconditioned skin (van den Broeke et al., 2016; 2017). Mechanical pinprick stimuli (64 and 96 mN) were applied before (“pre”) and 20 minutes after applying HFS (“post”) to the skin surrounding the site where HFS was delivered (“test area”) and to the corresponding skin area of the contralateral control arm. During pinprick stimulation both the pupil size and electroencephalogram were continuously recorded.

**Fig 1.**
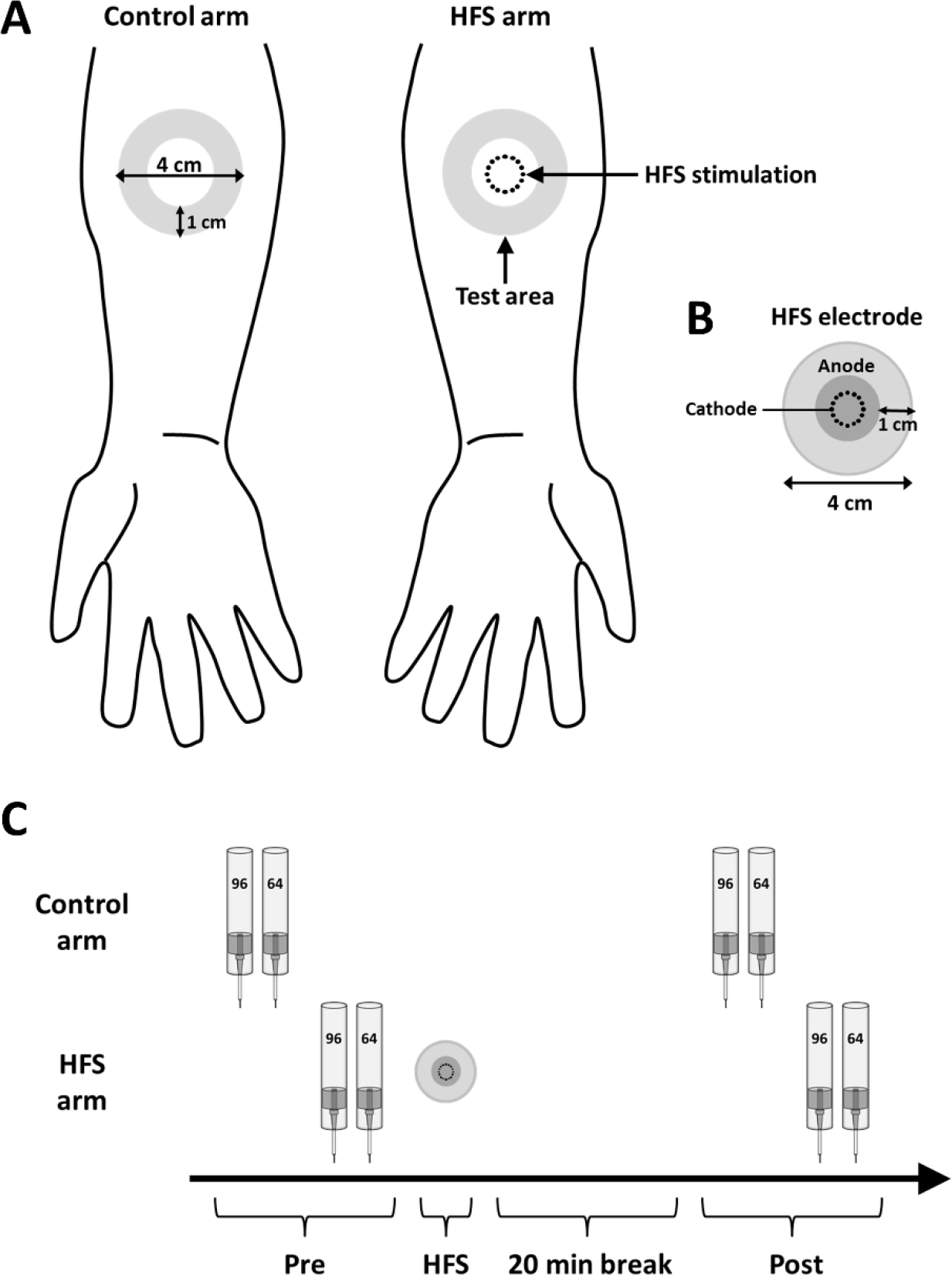
Experimental design. **A.** High frequency electrical stimulation of the skin (HFS) was applied to left or right volar forearm. Two different intensities of pinprick stimulation (64 mN and 96 mN) were applied to the skin surrounding the area onto which HFS was applied (“test area”) as well as to the same skin area on the contralateral control arm. **B.** Characteristics of the HFS electrode. **C.** Time-line of the experiment. The effect of pinprick stimulation on the perception, pupil size and brain responses was assessed at two different time-points: before HFS (“Pre”) and twenty minutes after applying HFS (“Post”).

The set-up of the experiment is shown in Figure 2A. The experiment was performed in a room with a constant and moderate level of ambient illumination. Participants were sitting in front of a table on an adjustable chair with their left or right arm inside a custom-built robotic pinprick stimulator (Fig. 2B). The head of the participant was rested on an adjustable chinrest which was mounted to the edge of the table. The chinrest was adjusted so that the gaze of the participant was at the middle of the computer screen, placed approximately 57 cm from the chinrest. The computer screen was used to display a fixation dot. An infrared camera, for recording pupil size, was placed in front of the computer screen approximately 35 cm from the chinrest.

**Fig. 2.**
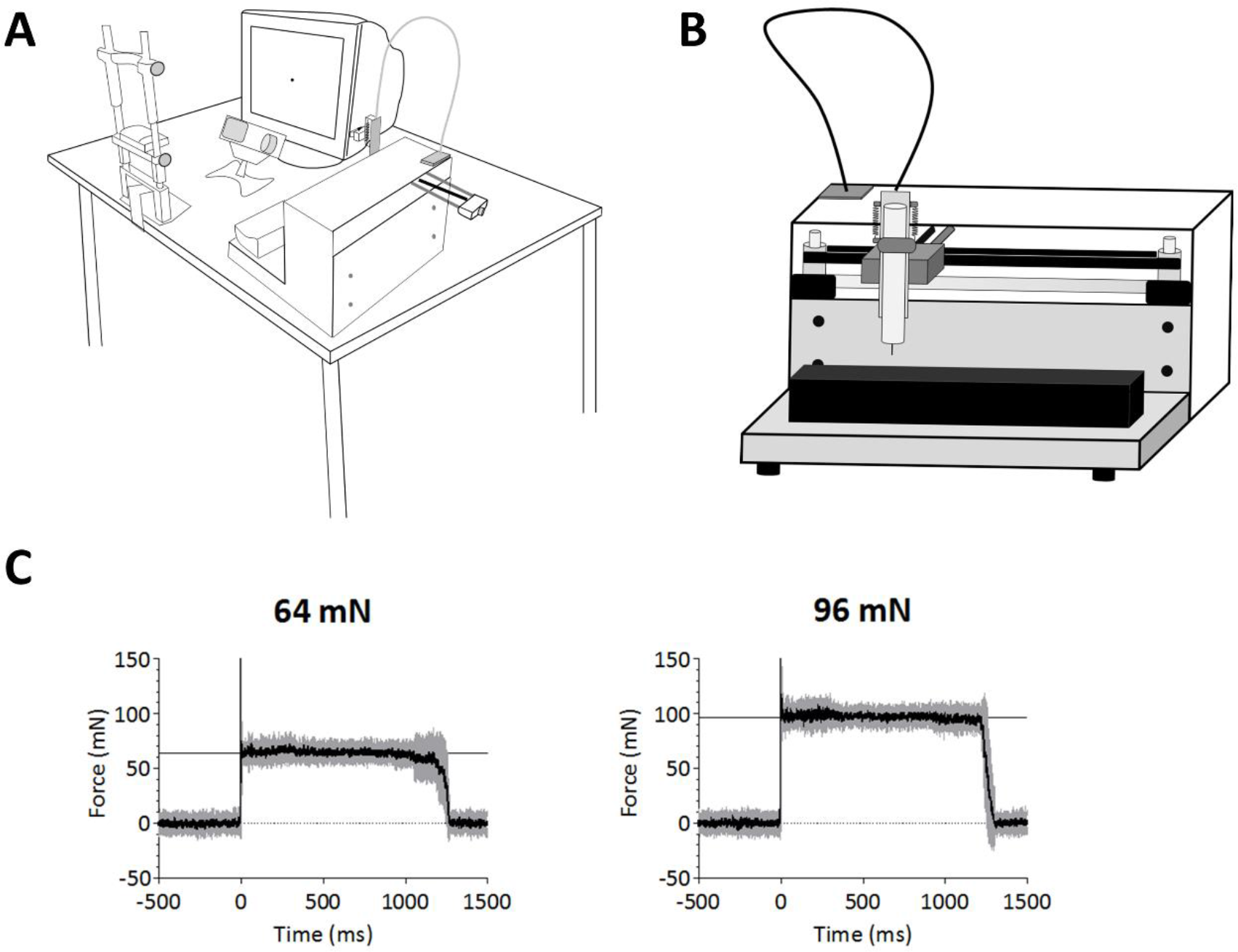
**A.** Experimental set-up. **B.** Schematic drawing of the robotic pinprick stimulator. **C.** Force profiles obtained during pinprick stimulation applied on a 6-axis strain-gauge force-torque transducer. Shown are the mean (black line) and standard deviation (grey lines) normal forces for both 64 mN and 96 mN pinprick stimulation. The mean was calculated across twenty trials.

#### High frequency electrical stimulation (HFS)

HFS consisted of five trains of 100 Hz electrical pulses (pulse width: 2 ms) lasting 1 s each. The time interval between the onsets of each train was 10 s. HFS was delivered to the volar forearm, 10 cm distal to the cubital fossa (Fig. 1A). The intensity of stimulation was individually adjusted to 20x the absolute detection threshold to a single pulse (0.34 ± 0.06 mA; mean ± sd). The electrical pulses were triggered by a programmable pulse generator (Master-8; AMPI Israel), produced by a constant current electrical stimulator (Digitimer DS7A, Digitimer UK), and delivered to the skin using a custom electrode (Fig. 1B) designed and built at the Centre for Sensory-Motor Interaction (Aalborg University, Denmark). The cathode consists of 16 blunt stainless-steel pins with a diameter of 0.2 mm protruding 1 mm from the base. The 16 pins are placed in a circle with a diameter of 10 mm. The anode consists of a surrounding stainless-steel ring having an inner diameter of 22 mm and an outer diameter of 40 mm. To avoid any confounding effect of handedness, the arm onto which HFS was applied (dominant vs. non-dominant) was counterbalanced across participants. Handedness was assessed using the Flinders Handedness Survey (Nicolls et al., 2011).

#### Robotic mechanical pinprick stimulation

A custom-built calibrated robot-controlled pinprick stimulator was used to deliver reproducible mechanical pinprick stimuli. The robot consists of three linear computer-controlled stages. The first two stages control the horizontal (X/Y) position of the pinprick probe. The third stage, onto which the pinprick probe is mounted, controls the vertical (Z) position of the probe. Each stage can be displaced at a speed of 25 mm/s, with a resolution of 0.1 mm. The pinprick probe consists of a stainless-steel flat tip probe (diameter: 0.35 mm, uniform geometry) on top of which rests a calibrated cylindrical weight. The probe and weight are mounted inside an aluminum tube, held by the robot. When applied perpendicular to the skin, the probe and weight slide freely inside the tube, thereby applying a constant normal force entirely determined by the total mass of the probe and weight. A high-resistance switch generates a trigger in the EEG that marks stimulation onset. When the probe touches the skin, this switch is triggered by the reduced impedance between the probe and an electrode placed on the skin at the wrist. A thin layer of conductive gel was applied onto the skin to lower the impedance between the contacting probe and the skin (van den Broeke et al., 2016; 2017). Before the start of each block, the X/Y/Z position of the pinprick stimulator was adjusted to position the probe approximately 5 mm above the skin, at the center of the test area. Two intensities of pinprick stimulation were used (64 and 96 mN). For each intensity of pinprick stimulation, each arm (HFS and control arm), and each time point (pre and post), a total of twenty stimuli were administered, in separate blocks. The time-course of the stimulus is shown in Figure 2C. The order of presentation of the two pinprick intensities, as well as the arm onto which stimuli were first applied (HFS vs. control arm) were counterbalanced across participants. The pinprick robot stimulated at random positions within the test area and never the same spot twice by first displacing the probe to the corresponding position in the X/Y plane, and the performing a downward 10 mm movement (Z-axis). After 1.2 s, the pinprick probe was removed by performing an upward 10 mm movement. Both movements were performed at a constant speed of 16.66 mm/s. The test area was defined at the beginning of the experiment by drawing a circle of 4 cm diameter that exactly matches the size of the HFS electrode at the ventral forearm (Fig. 1). During the pinprick stimulation, a 48 × 57.5 cm panel with an opening for the arm was placed in front of the pinprick robot to prevent view of the stimulated arm. Moreover, during pinprick stimulation, participants listened to white noise via headphones to mask any sound generated by the pinprick robot during movements.

Each trial started with the presentation of the fixation dot on the computer screen, which remained during the whole trial. After the fixation dot appeared, the pinprick stimulus was delivered onto the skin after a randomized interval between 2 to 4 seconds. Five seconds after delivery of the pinprick stimulus, the fixation dot disappeared, indicating the end of the trial. The next trial started after an inter-stimulus interval (ISI) of 2 seconds. During each trial, participants were instructed to fixate their gaze at the dot and to avoid blinking and moving. Participants were instructed to blink after the dot disappeared and until the next trial started, indicated by the re-appearance of the fixation dot.

#### Intensity of perception

To confirm the successful induction of increased pinprick sensitivity at the HFS-stimulated arm, participants were asked to report, after each block, the average intensity of perception of the pinprick stimuli using a numerical rating scale (NRS) ranging from 0 (no perception) to 100 (maximal pain), with 50 representing the transition from non-painful to painful domains of sensation. To assess changes in intensity of perception, we performed for both stimulation intensities (64 and 96 mN) a General Linear Model (GLM) repeated measures ANOVA using two within-subject factors: time (pre vs. post) and arm (control vs. HFS arm). The dependent variable was the NRS score. The level of significance was set at p<.05. For post-hoc testing we compared the effect of time separately for each arm (control and HFS), therefore, the p-value was Bonferroni corrected (p<.025). Effect sizes were calculated using the classical Cohen’s *d*. The statistical analyses were conducted using SPSS 18 (SPSS Inc., Chicago, IL, USA).

#### Pupil size recording

The pupil diameter of the right eye was recorded using an infrared eye-tracker (EyeLinks 1000; SR Research Ltd, Kanata, ON, Canada) at a sampling rate of 1000 Hz. As is usual with eye-tracking and pupillometry experiments, the eye was calibrated using a nine-point calibration and validation procedure (e.g. Mathôt, Dalmaijer, Grainger and Van der Stigchel, 2014). The pupil diameter was recorded whilst subjects fixated on a small black fixation dot which was presented on a 19” viewsonic G90fB Graphics series monitor, set at a 1024 × 768 resolution and a 60 Hz refresh rate. On screen stimuli and communication with the eye-tracker were generated and displayed using a custom coded MATLAB script and a set of procedures allowing precise timing of the display and synchronization with the eye-tracker (Brainard, 1997; Cornelissen et al., 2002; Pelli, 1997). Calibration was considered successful when the error was <1° of visual angle.

#### EEG recording

The EEG was recorded using 32 actively shielded Ag-AgCl electrodes that were mounted in an elastic cap and arranged according to the international 10-20 system (Waveguard32 cap, Advanced Neuro Technologies, The Netherlands). The EEG signals were amplified and digitized using a sampling rate of 1000 Hz and an average reference (HS32; Advanced Neuro Technologies, The Netherlands). Eye movements were recorded using two surface electrodes placed at the upper-left and lower-right sides of the left eye. Electrode impedances were kept below 20 kΩ.

#### Pupil size analyses

The continuous signals expressing pupil diameter as a function of time were preprocessed using Matlab (Matlab R2016, The MathWorks Inc., Natick, Massachusetts, United States) and further analyzed using the Letswave 6 toolbox (nocions.org/letswave). The signals were segmented into epochs according to the beginning and end of each trial. Time intervals during which pupil diameter was missing because of occasional eye blinks were linearly interpolated. The data were then further segmented into epochs extending from -2000 to +6200 ms relative to the onset of pinprick stimulation and a baseline correction was applied (reference interval ranging from -2000 to 0 ms). Finally, separate average waveforms were computed for each participant, time point (pre and post), stimulation site (HFS and control) and stimulation intensity (64 and 96 mN).

To assess whether there was a significant difference after HFS between the pupil size waveforms elicited from both stimulation sites, we used a non-parametric cluster-based permutation approach (van den Broeke et al., 2017). To summarize, we first computed for each subject difference waveforms assessing the change in pupil size at post vs. pre at the control arm (control arm_post_ – control arm_pre_) and at the HFS-treated arm (HFS arm_post_ – HFS arm_pre_). Then, we applied the cluster-based permutation test on the difference waveforms of both arms. The test, adapted from Van den Broeke et al. (2017) and summarized here, consisted of the following steps. First, the difference waveforms were compared by means of a point-by-point paired-sample t-test. Then, samples above the critical t-value for a parametric two-sided test that were adjacent in time were identified and clustered. An estimate of the magnitude of each cluster was then obtained by computing the sum of the absolute t-values constituting each cluster (cluster-level statistic). Random permutation testing (2000 times) of the subject-specific difference waveform of the two arms (performed independently for every subject) was then used to obtain a reference distribution of maximum cluster magnitude. Finally, the proportion of random partitions that resulted in a larger cluster-level statistic than the observed one (i.e. p-value) was calculated. Clusters in the observed data were regarded as significant if they had a magnitude exceeding the threshold of the 95th percentile of the permutation distribution, which corresponds to a critical alpha-level of 0.05.

To assess changes in baseline pupil size, a GLM repeated measures ANOVA was performed on the average pupil size over the first 2 seconds of the prestimulus period using three within-subject factors: time (pre vs. post), arm (control vs. HFS arm) and intensity (64 mN vs. 96 mN).

#### EEG analysis

The EEG signals were analyzed offline using the Letswave 6.0 toolbox for Matlab.

### Pinprick-evoked potentials (PEPs)

The continuous EEG signals were bandpass-filtered using 0.3-30 Hz band pass zero-phase Butterworth filter. The signals were then segmented into epochs extending from -500 to +1500 ms relative to stimulus onset. Epochs contaminated by eye movements or eye blinks were corrected using an Independent Component Analysis (ICA; (Jung et al., 2000)). Denoised epochs were then baseline-corrected (reference interval: -500 to 0 ms) and re-referenced to linked earlobes (A1A2). Finally, epochs with amplitude values exceeding ±100 μV were rejected as these were likely to be contaminated by artifacts. Separate average waveforms were computed for each participant, time point (pre and post), stimulation site (HFS and control) and stimulation intensity (64 and 96 mN). To assess whether there was a significant difference after HFS between the PEP waveforms elicited from both stimulation sites, we used the same approach as for analyzing pupil size. We first computed for each subject difference waveforms (post vs. pre) for both arms separately and then we applied the non-parametric cluster-based permutation test on these difference waveforms. Based on the results of our previous studies (van den Broeke et al., 2015; 2016; 2017) we performed the permutation test only on the midline central-posterior electrodes (i.e. electrode Cz and Pz). Because we performed the permutation test on two electrodes we have set the critical p-value at p<.025 (.05/2).

### Time-frequency analysis

We previously showed that the effects of HFS on pinprick-evoked EEG can also be identified by characterizing the change in EEG amplitude in the time-frequency domain (van den Broeke et al., 2017). The time-frequency analysis was performed as follows. First, we applied a short-time fast Fourier transform (STFFT) with a fixed Hanning window of 500 ms to the band-pass filtered (0.3-30 Hz) and ICA-denoised EEG signals used for the analysis of the PEPs. The explored frequencies ranged from 1 to 30 Hz. The STFFT was applied to each single EEG epoch. Then, the single-trial time-frequency estimates of oscillation amplitude were averaged across trials, separately for each subject and condition. To test for significant differences in the time-frequency representations after HFS between the two arms we analyzed the time-frequency maps obtained at electrodes Cz and Pz in the same way as for the pupil and PEP waveforms. Because our previous study revealed only significant changes in low-frequency EEG activity between 150-400 ms (van den Broeke et al., 2017), we restricted the cluster-based permutation test to the frequencies 1-7 Hz and the time points 0-500 ms. The critical p-value was set at p<.025. To visualize changes in post-stimulus EEG activity relative to the pre-stimulus baseline period, a baseline correction was applied. For each estimated frequency line, using the average signal amplitude between -500 to -100 ms relative to stimulus onset was subtracted from each post-stimulus time point.

## RESULTS

### Intensity of perception

HFS induced a clear increase in pinprick sensitivity at the HFS arm, as shown by the changes in the intensity of perception elicited by both 64 mN and 96 mN pinprick stimulation (Fig. 3A). This was confirmed by the repeated-measures ANOVA, which showed a significant time × treatment interaction for both the 64 mN stimulus (F(1,13)=61.467, p<.001) and the 96 mN stimulus (F(1,13)=43.216, p<.001). Post-hoc tests showed, for both intensities, a statistically significant increase of the perceived intensity at the HFS-treated arm after HFS (64 mN: paired t-test; t(13)=7.653, p<.0001, Cohen’s *d*=2.182; 96 mN: t(13)=6.206, p<.0001, Cohen’s *d*= 1.838). No significant changes in perceived intensity were observed at the control arm. Figure 3B shows, for both pinprick intensities, the increase in intensity of perception compared to baseline and control site.

**Fig. 3.**
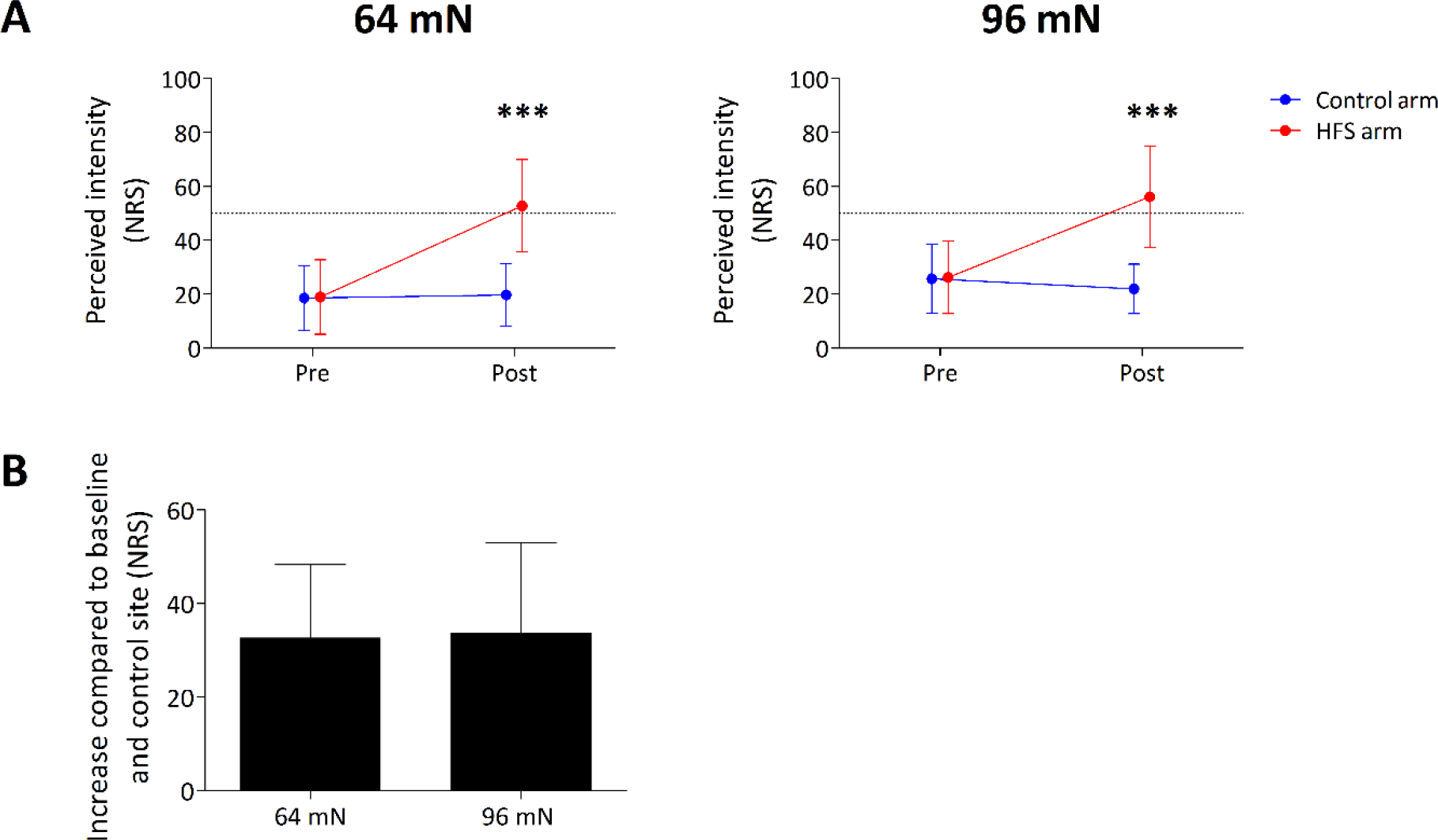
**A.** Intensity of perception elicited by the mechanical pinprick stimulation before and twenty minutes after applying HFS. Shown are the group-level average and standard deviation of the numerical rating scale (NRS) scores for the 64 mN and 96 mN stimulation intensity. Asterisks denote a statistically significant increase of the NRS scores after HFS at the HFS arm. **(B)** Group-level average and standard deviation increase in NRS compared to baseline and control site for the 64 mN and 96 mN stimulation intensity.

### Pupil size

The group-level average waveforms of pupil size are shown in Figure 4. Both before and after HFS, the 64 mN and 96 mN pinprick stimulus elicited an increase in pupil size. This pupil dilation response was increased after HFS at the HFS-treated arm. This was confirmed by the cluster-based permutation test which revealed a significant difference between the subtracted waveforms (post-pre) of both arms (control vs. HFS) for both the 64 mN stimulus (p<.001) and the 96 mN stimulus (p<.01). Post-hoc tests performed on the individual magnitudes of post-stimulus activity (average pupil size between 0-5 s) showed, for both intensities, a statistically significant increase of the pupil size at the HFS-treated arm after HFS (64 mN: paired t-test; t(13)=4.452, p<.001, Cohen’s *d*= 1.288; 96 mN: t(13)=2.603, p<.05, Cohen’s *d*= 0.687). No significant changes in pupil size were observed after HFS at the control arm. Figure 5 shows, for both pinprick intensities, the changes in pupil size compared to baseline for each subject at both arms. The increase in pupil size was, on average, greater for the 64 mN stimulus as compared to the 96 mN stimulus, and this difference was marginally significant (p=.0503, Fig. 5B). Figure 6 shows the group-level average (and sd) pupil size of the first two seconds of the pre-stimulus baseline interval. The repeated measures ANOVA did not show a significant *time × side × intensity* interaction (F(1,13)=.259, p=.619) or significant *time × intensity* interaction (F(1,13)=1.667, p=.219) on the magnitude of the baseline pupil size.

**Fig. 4.**
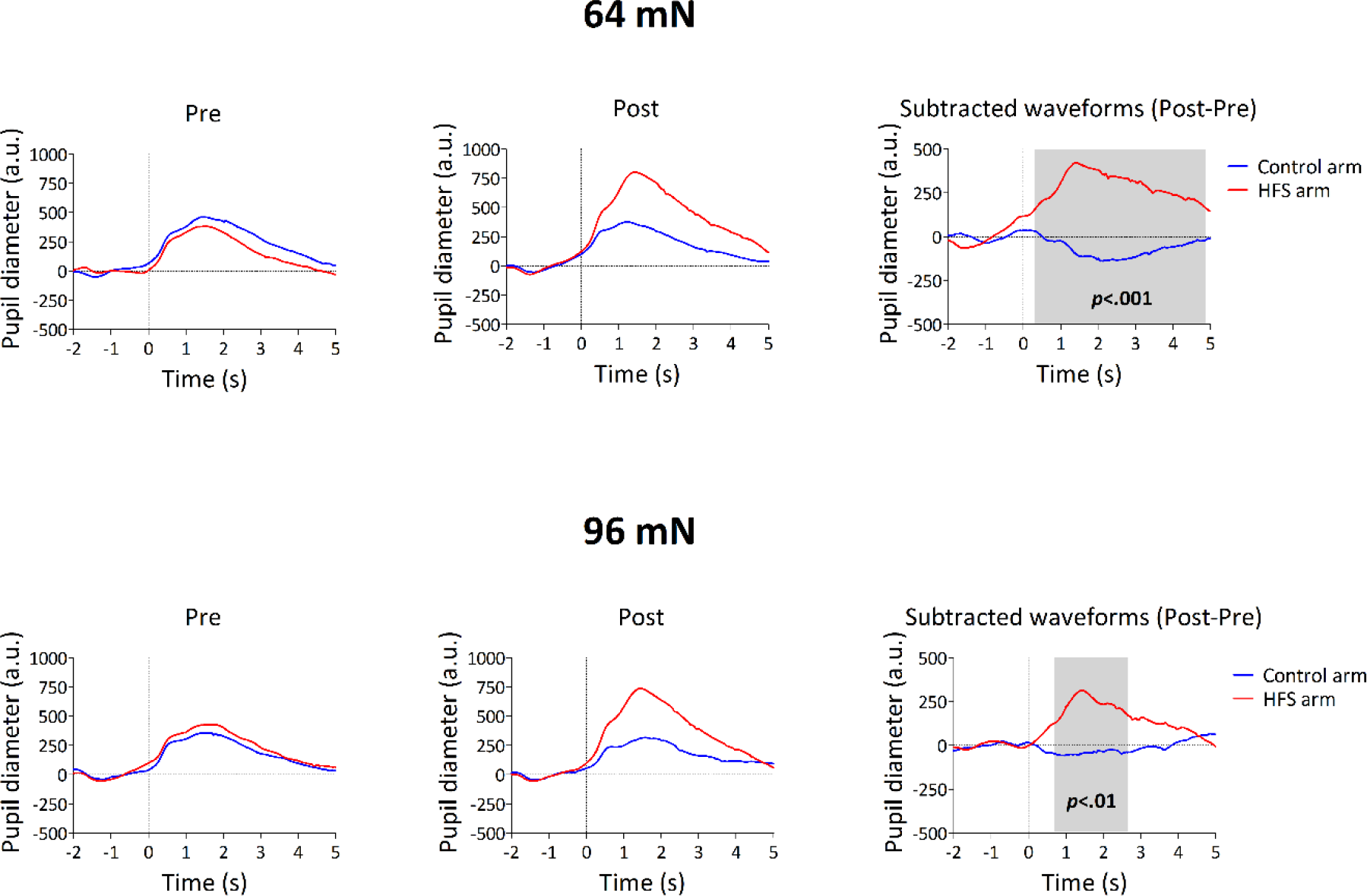
Pupil responses elicited by the mechanical pinprick stimulation before and after HFS. Shown are the group-level average pupil waveforms for both arms and both stimulation intensities, as well as the subtracted waveforms (post minus pre). The grey rectangle indicates the time window at which the two waveforms were statistically significant different. X-axis shows the time (in seconds) relative to stimulus onset, whereas the Y-axis shows in arbitrary unites the size of the pupil. Positive values refer to increases in pupil size whereas negative values refer to decrease in pupil size.

**Fig. 5.**
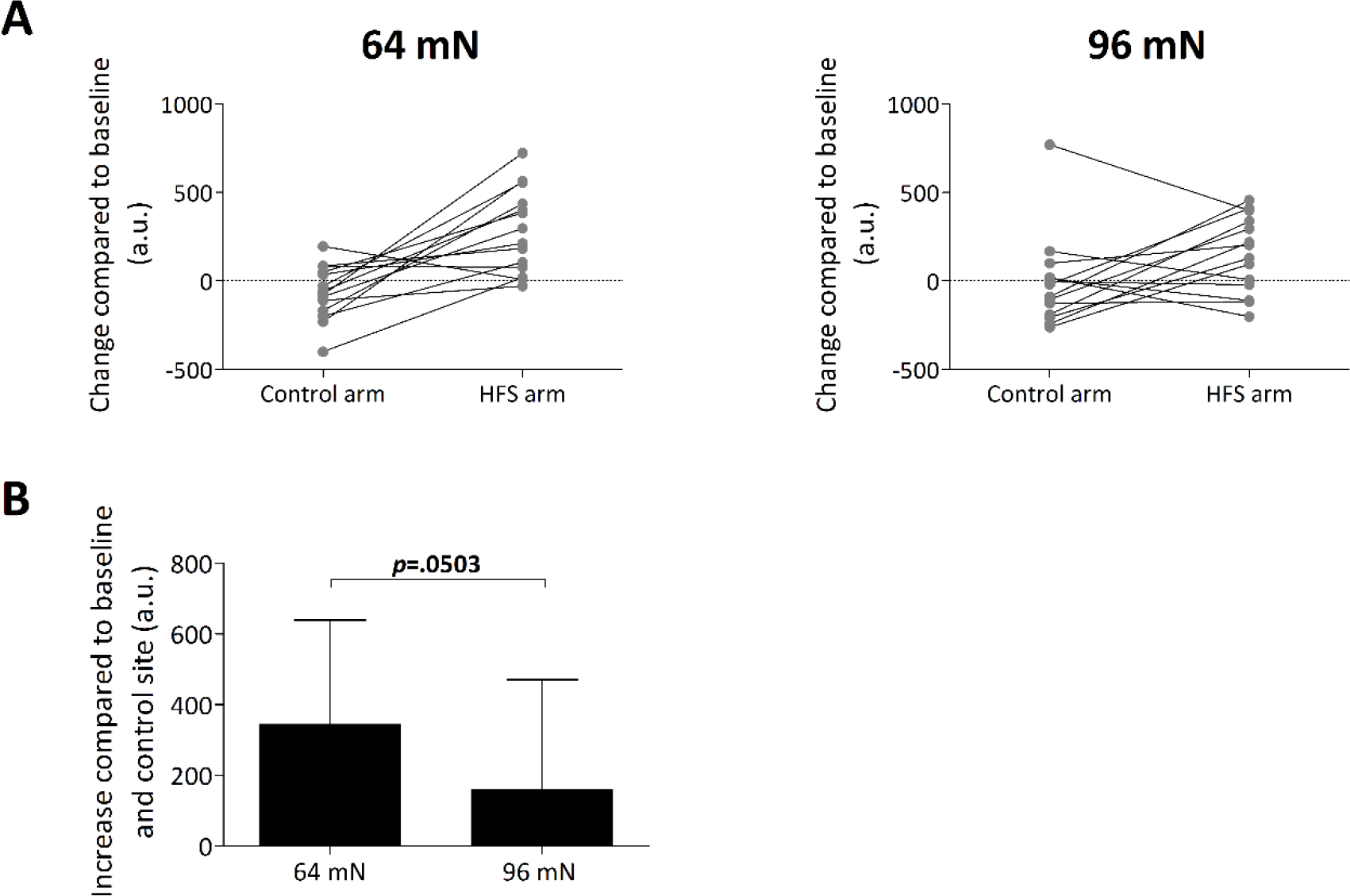
**A.** Individual magnitudes of the change in pupil size relative to baseline for both the 64 and 96 mN stimulation intensity. **B.** Group-level average and standard deviation increase in pupil compared to baseline and control site for the 64 and 96 mN pinprick intensity.

**Fig. 6.**
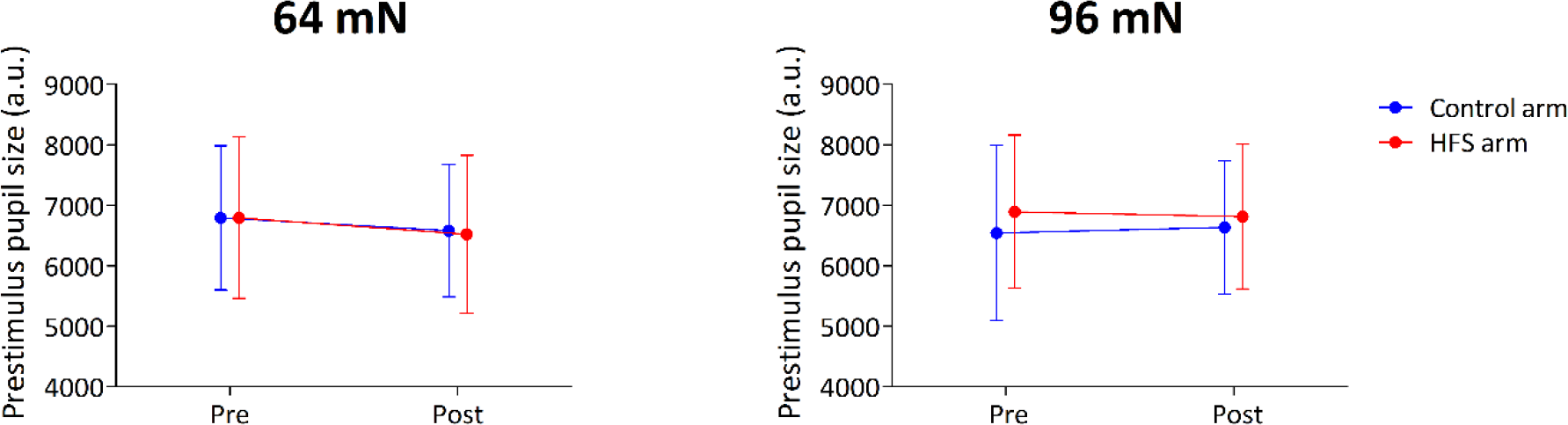
Group-level average and standard deviation of the baseline pupil size (arbitrary unites) before and after HFS for both arms and both stimulation intensities.

### Pinprick-evoked brain potentials (PEPs)

The group-level average waveforms of PEPs elicited by the 64 mN and 96 mN stimulation are shown in Figure 7. For both stimulation conditions, the waveforms appeared as an early-latency negative wave, peaking approximately 120 ms after stimulation onset, followed by a later positive wave, peaking approximately 320 ms after stimulation onset. Both waves were maximal at the scalp vertex. As compared to our previous study (van den Broeke et al., 2016), the negative wave was markedly more visible. This could be related to the improved reproducibility of the pinprick stimulation (manual versus robot). The cluster-based permutation test did not reveal a statistically significant difference between the subtracted waveforms (post-pre) of both arms (control vs. HFS) for neither the 64 mN nor 96 mN pinprick stimulation.

**Fig. 7.**
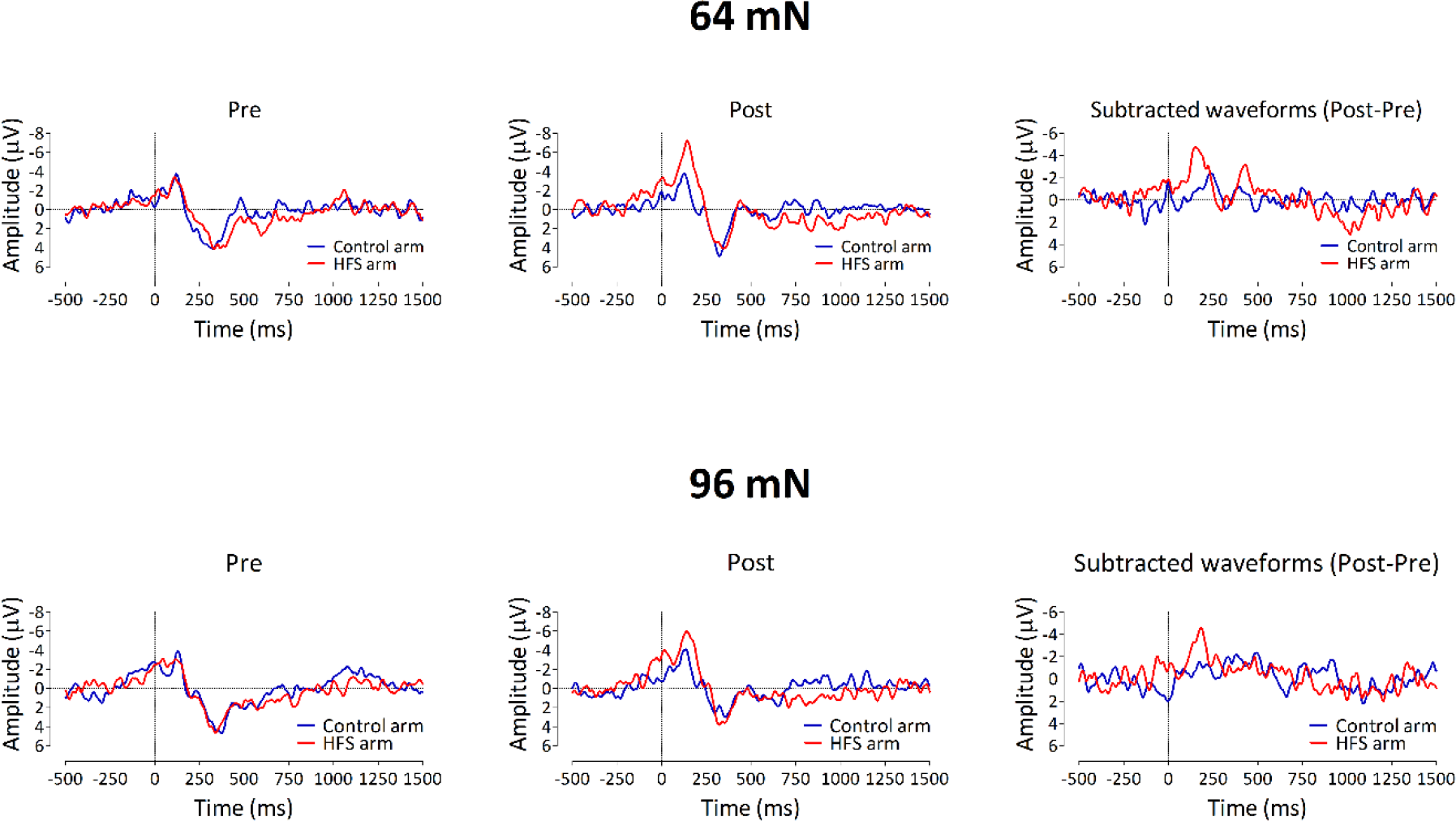
Pinprick-evoked brain responses (PEPs) elicited by mechanical pinprick stimulation before and after HFS. Shown are the group-level average PEP waveforms for both arms and both stimulation intensities, as well as the subtracted waveforms (post minus pre). X-axis shows the time (in milliseconds) relative to stimulus onset, whereas the Y-axis shows the amplitude of the responses. Shown are the PEP waveforms from Cz vs. A1A2.

### Time-frequency analysis of low-frequency EEG activities (1-7 Hz)

The group-level average time-frequency maps obtained at both time points (pre and post HFS) and arms (control and HFS) are shown in Figure 8. For the 64 mN stimulus the cluster-based permutation test performed on the difference time-frequency maps (post vs. pre HFS) between both arms (control vs. HFS) revealed a marginally significant cluster between 100 and 250 ms and between 3 and 7 Hz (p=0.0297, Fig. 8B). No significant clusters were identified for the 96 mN stimulus.

**Fig. 8.**
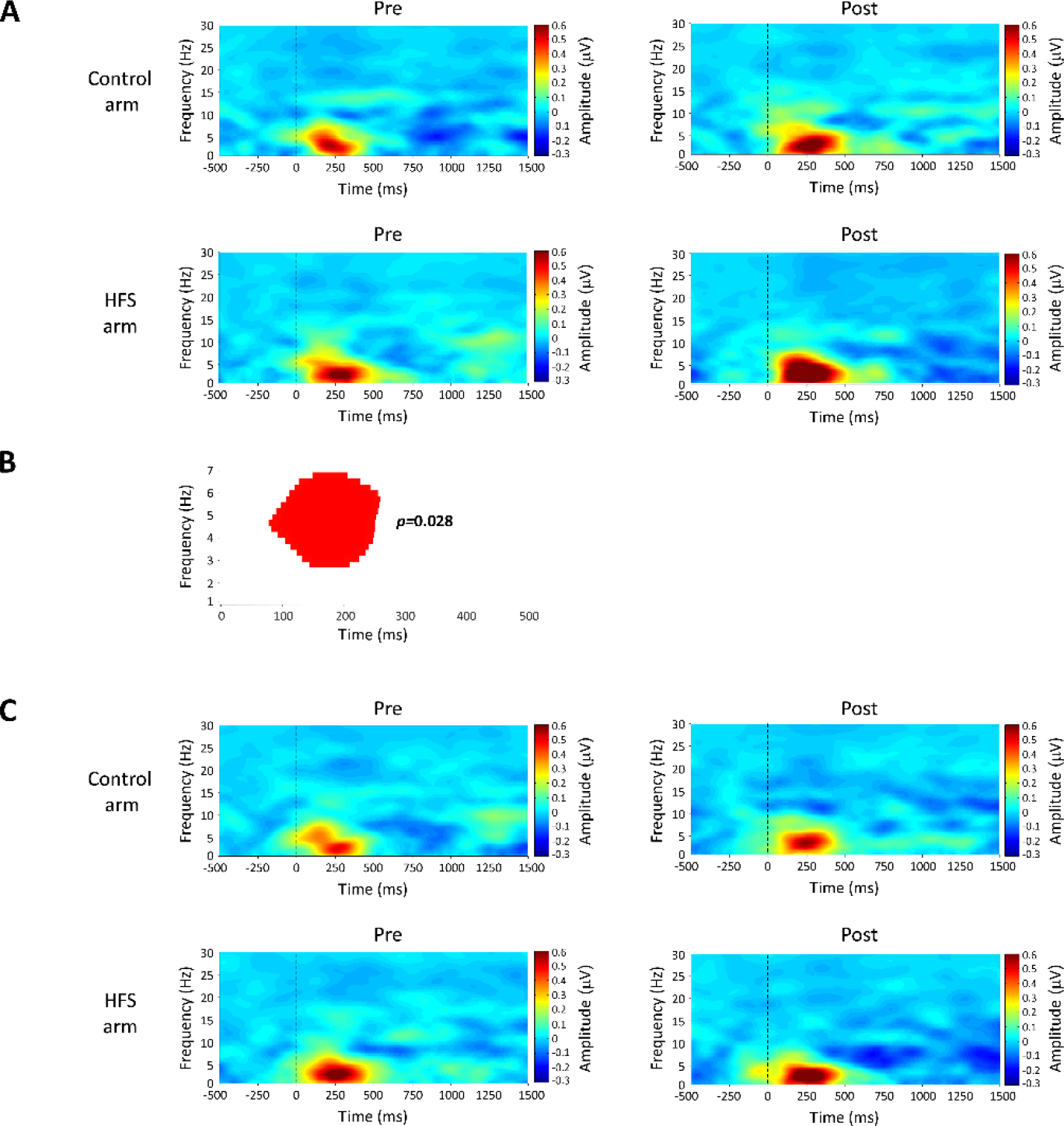
Time-frequency representations (TFR) of the EEG responses elicited by the pinprick stimulation before and after HFS. **A.** Group-level average TFRs for the 64 mN stimulation intensity. X-axis shows the time (in milliseconds) relative to stimulus onset, whereas the Y-axis shows the range of frequencies. The red colour represents an increase in amplitude whereas the blue colour represents a decrease in amplitude. TFRs are recorded from Cz vs. A1A2. **B.** Cluster of time and frequency points at which the cluster-based permutation test identified a marginally significant interaction in the EEG responses elicited by the 64 mN stimulation intensity. **C.** Group-level average TFRs for the 96 mN stimulation intensity. Shown are the TFRs from electrode Cz vs. A1A2.

### Post-hoc analyses

#### Negative wave of the PEPs

Although the magnitude of the pinprick-evoked negative wave elicited by pinprick stimulation of the HFS arm appeared to be, on average, greater after HFS as compared to before HFS, especially for the 64 mN stimulus, the cluster-based permutation tests revealed no significant cluster. However, the time-frequency analysis revealed a marginally-significant interaction whose latency corresponded to the latency of this negative wave. Moreover, the non-parametric cluster-based permutation approach may be less sensitive when the PEP responses are limited to a small number of time points and show great jitter across participants. For these reasons, we determined individual peak values of the N wave (maximal negative peak within the arbitrary time window 80-180 ms), to test if the N wave is significantly enhanced after HFS at the HFS-treated arm. Such as for the analysis of NRS scores, we performed, for both stimulation intensities (64 and 96 mN), a repeated measures ANOVA using time (pre vs. post) and arm (control vs. HFS) as within-subject factors. The RM-ANOVA showed a trend towards a significant interaction (F(1,13)=4.187, p=.062) for the N-wave elicited by the 64 mN intensity. The effect size of the increase in N wave at the HFS-treated arm after HFS, Cohen’s *d*= 0.833, suggests that the lack of significant interaction might be due to the small sample size. The sample size of the present study was based on a sample size calculation performed on the results of the previous study (van den Broeke et al., 2017) which indicated that with such an effect size and within-subject design a total of eleven subjects would be needed to detect an increase in the late positive wave of PEPs elicited by the 64 mN pinprick stimulus.

No significant interaction or trend towards a significant interaction was observed for the N-wave elicited by the 96 mN intensity (F(1,13)=1.784, p=.205).

#### Absence of increase in the positive wave of the PEPs after HFS

The cluster-based permutation test did not reveal a significant difference between the subtracted pinprick-evoked waveforms of both arms for the 64 mN pinprick stimulation. In order to estimate the effect size we calculated individual peak values of the positive wave which were determined as the maximal positive peak within the arbitrary time window 250-500 ms. The observed effect size was 0.040 (Cohen’s *d*), which strongly indicates that there was no effect of HFS on the positive wave of the PEPs.

The changes in magnitude of the N- and P-waves after HFS (compared to baseline) elicited by both pinprick intensities are shown in Fig. 9. Figure 10 shows the increase in N- and P-wave compared to baseline and control site.

**Fig. 9.**
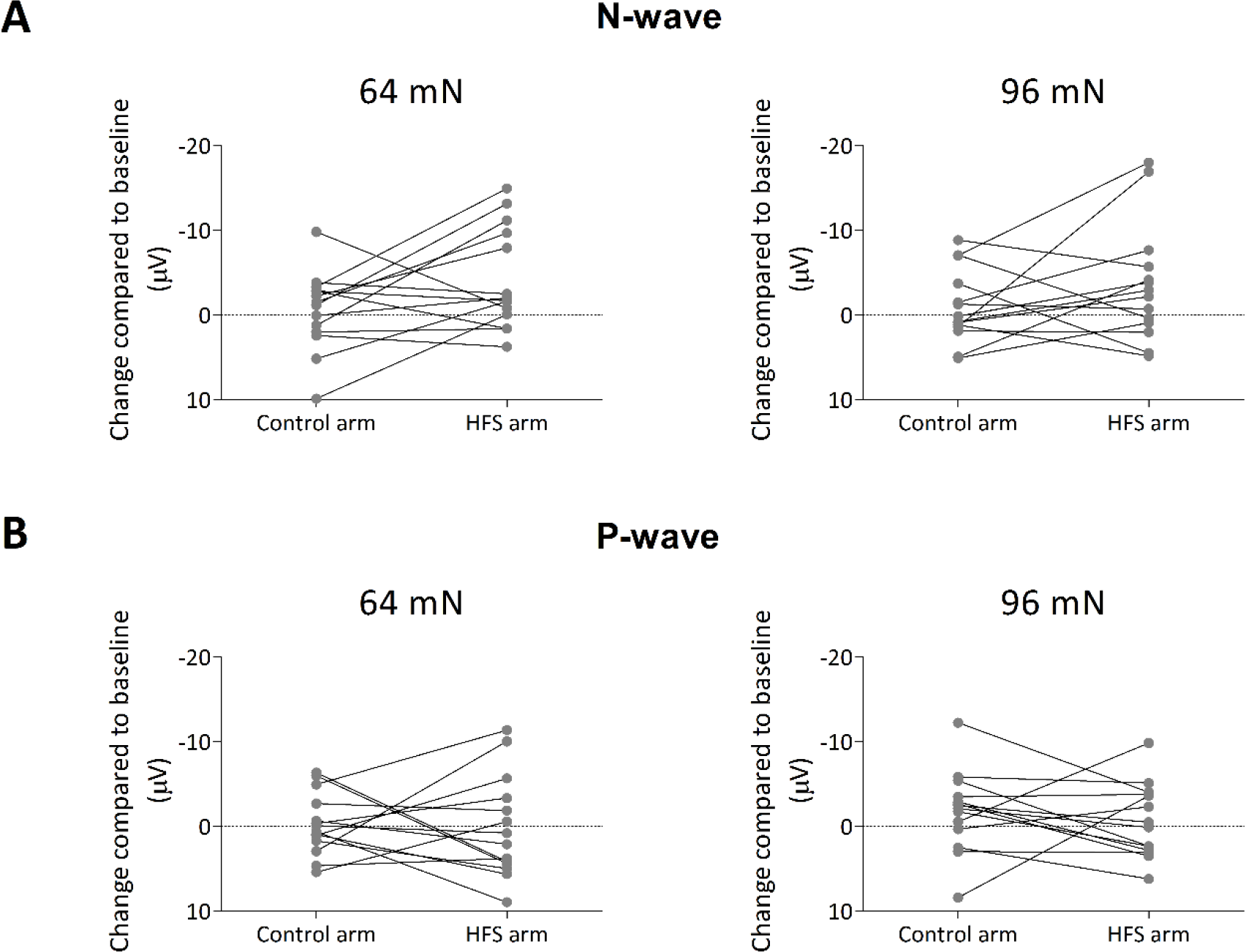
Individual magnitudes of the change in N-wave **(A)** and P-wave **(B)** of the PEPs compared to baseline for both the 64 mN and 96 mN stimulation intensity.

**Fig. 10.**
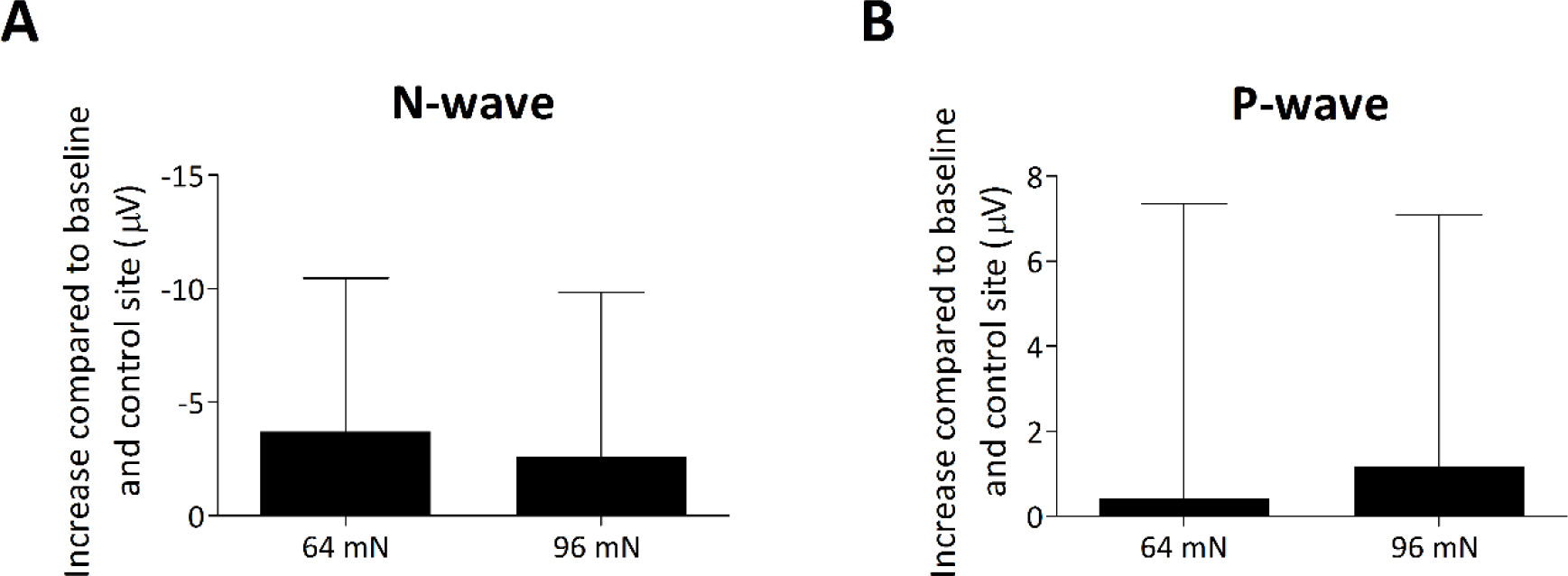
Group-level average and standard deviation increase in N-wave **(A)** and P-wave **(B)** of the PEPs compared to baseline and control site for the 64 mN and 96 mN stimulation intensity.

## DISCUSSION

In the present study we show that mechanical pinprick stimuli delivered to the area of increased pinprick sensitivity elicits a pupil dilation response of greater amplitude as compared to mechanical pinprick stimuli delivered before HFS or to control site. However, contrary to our previous study (van den Broeke et al., 2017), no significant increase in the magnitude of the late positive wave of PEPs was observed after HFS.

### HFS increases the pinprick-evoked pupil dilation response

HFS significantly increased the pupil dilation response elicited by pinprick stimuli delivered to the HFS-treated arm. This was the case for both pinprick intensities, even though the increase tended to be larger for the 64 mN stimulus as compared to the 96 mN stimulus. This seems to be in line with our previous results (van den Broeke et al., 2017) showing a greater enhancement of the late positive wave of PEPs elicited by the 64 mN stimulation as compared to 96 mN stimulation. Taken together, this suggests a dissociation between pinprick perception (which tends to be similarly enhanced for both pinprick intensities) and both the pupil response and PEPs (which tend to be preferentially enhanced for intermediate intensities of pinprick stimulation, 64 mN).

No significant difference in the baseline pupil diameter was observed after HFS between both pinprick intensities. Hence, our study does not provide evidence that, after HFS, baseline LC activity is higher during the 96 mN stimulation of the area showing increased pinprick sensitivity compared to the 64 mN stimulation.

Recently, Joshi et al. (2016) showed that direct electrical stimulation of LC neurons triggered pupil dilations. However, at present there are no known direct connections between LC and brainstem nuclei controlling pupil size, which raises the question of how the two are related (Joshi et al., 2016; Costa and Rudebeck, 2016). Regarding pupil dilations triggered by external events, one of the possibilities that has been suggested is that the LC and the nuclei controlling pupil size are co-modulated by a third region, of which the nucleus paragigantocellularis (PGi) of the ventral medulla is a possible candidate (Nieuwenhuis et al., 2011; Joshi et al., 2016; Costa and Rudebeck, 2016). To the best of our knowledge it is currently unknown whether mechanical pinprick stimuli activate the PGi, however, a previous study showed that PGi neurons projecting to the LC were excited by hindpaw stimulation (Ennis and Aston-Jones, 1987). Moreover, the pharmacological blockade of PGi-LC excitatory pathways also blocks the LC responses to sciatic nerve stimulation (Ennes and Aston-Jones, 1988). Taken together, mechanical nociceptive input from the dorsal horn may activate the PGi in the brainstem that elicits activity in LC neurons in parallel with autonomic reflex responses (Sara and Bouret, 2012).

### Lack of increase of the late positive wave of PEPs after HFS

Despite the clear increase in pinprick perception and pupil size after HFS, we did not observe any significant increase in the magnitude of the late positive wave of PEPs after HFS at the HFS treated arm. The marked difference in effect sizes suggests that the lack of a significant effect of HFS on the positive wave of PEPs vs. the presence of a significant effect of HFS on pupil size was truly due to a differential effect of HFS on these two outcome measures. Importantly, the lack of effect of HFS on the late positive wave of PEPs is in striking contrast with our previous study (van den Broeke et al., 2017). Note that in that previous study the effect size (Cohen’s *d*) of the increase in positive wave at the HFS-treated arm after HFS was 1.018.

One important difference between the present study and that previous study is the task participants had to perform *during* pinprick stimulation. In the previous study, participants had to report the quality of the percept elicited by each pinprick stimulus. In addition, they had to provide a numeric rating scale score of the intensity of perception at ten pseudo-randomly selected trials of block of thirty trials. Hence, the pinprick stimuli were task relevant. In the present study, participants were instructed to provide an average rating of the pinprick stimuli only at the end of each block. Hence, the stimuli were probably less task relevant. This task instruction had been chosen because we wanted to avoid that providing a response directly after each pinprick stimulus would influence the pupil response.

It could thus be that the lack of effect of HFS on the late positive wave of PEPs in the present study was due to the low task relevance of the pinprick stimulus. Indeed, the late vertex positivity elicited by nociceptive laser stimuli has been shown to be strongly dependent on task-relevance and/or the amount of attention allocated to the stimulus (Siedenberg and Treede, 1996; Legrain et al., 2002; 2003).

Therefore, it is quite possible that the increase in magnitude of the positive wave of PEPs that can be induced by HFS is strongly dependent on the task and/or the active engagement of the participant. In contrast, the HFS-induced increase in pinprick-evoked pupil dilation seems not or less affected by these task requirements.

Interestingly, the magnitude of the negative wave elicited by the 64 mN pinprick intensity tended to be increased after HFS at the HFS-treated arm. Furthermore, this did not appear to be so much the case for the negative wave of the PEPs elicited by the 96 mN stimulus. This was similar to the increase in the pupil response, which also tended to be greater for the 64 mN stimulus as compared to the 96 mN stimulus, suggesting that both responses may at least partly depend on shared mechanisms.

To conclude, in the present study, we observed that HFS enhances the magnitude of the pupil response elicited by pinprick stimulation of the skin showing increased pinprick sensitivity. The finding that this enhancement tended to be greater at 64 mN stimulation as compared to 96 mN stimulation is in line with our previous results also showing a stronger enhancement of PEPs at 64 mN compared to 96 mN stimulation (Fig. 11). Assuming that the pupil response reflects phasic LC activity, these observations are compatible with the hypothesis that PEPs relate, at least in part, to phasic LC activity.

**Fig. 11.**
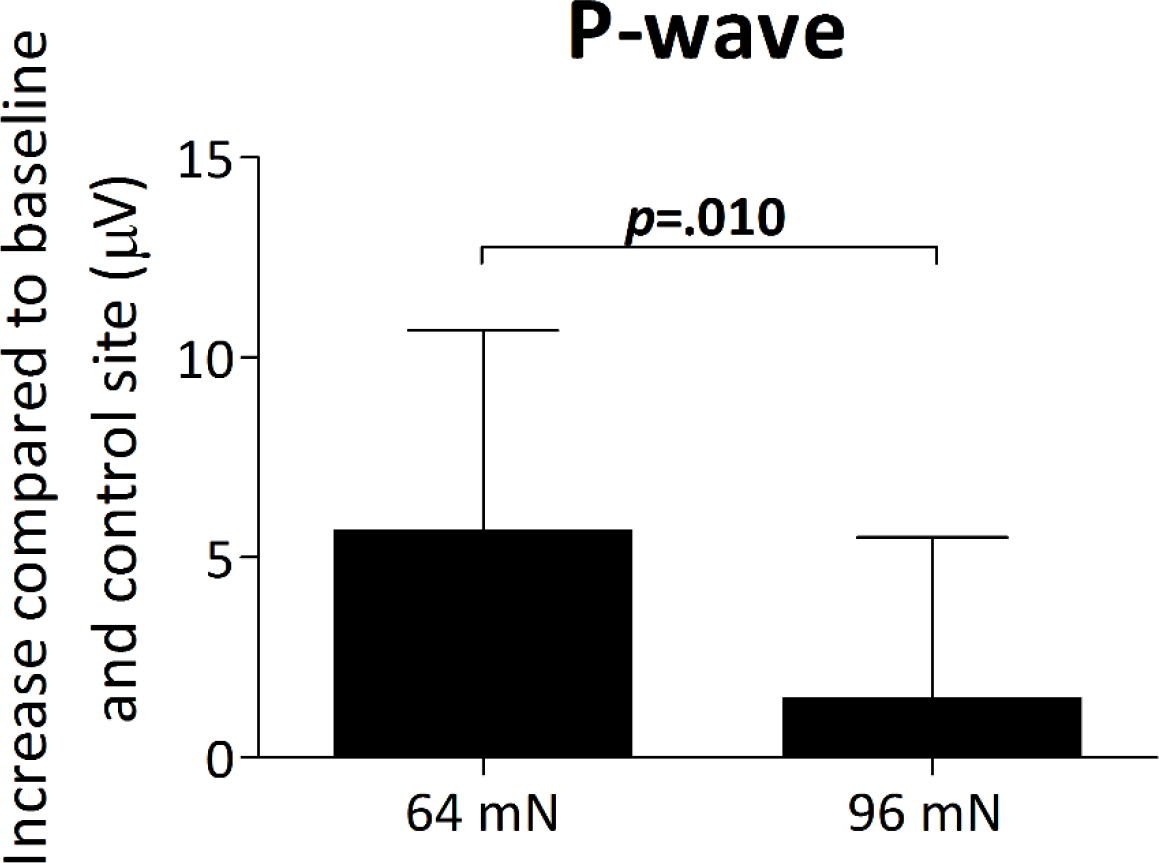
Group-level average and standard deviation increase in positive wave of the PEPs compared to baseline and control site for the 64 and 96 mN stimulation intensity as observed in our previous study (van den Broeke et al., 2017). The p-value refers to the results of a paired t-test.

However, the lack of increase in the positive wave of PEPs in the present study, suggest that its occurrence is also strongly dependent on additional factors, which could be task relevance or attentional engagement. Future studies should further investigate this. Most importantly, pupil size appears to be a more sensitive measure for detecting the effects of central sensitization than PEPs.

## CONFLICT OF INTEREST

None declared.

### AUTHOR CONTRIBUTION

EvdB and AM contributed to the conception and design of the experiment. EvdB, MH and JB contributed to the data acquisition. EvdB and JL performed the analyses. EvdB and AM contributed to the interpretation of the data. EvdB, MH, JB, JL and AM approved the final version of the manuscript.

### FUNDING

EvdB, JL and AM are supported by the ERC “Starting Grant” (PROBING PAIN 336130). EvdB is also supported by the Fonds de Recherche Clinique (FRC) provided by the Université catholique de Louvain (UCL), Belgium. MH is supported by a travel grant provided by the Behavioral Science Institute (BSI) of the Radboud University, The Netherlands. JB is supported by a grant provided by the Université catholique de Louvain (UCL), Belgium.

